# cytoNet: Spatiotemporal Network Analysis of Cell Communities

**DOI:** 10.1101/180273

**Authors:** Arun S. Mahadevan, Byron L. Long, Chenyue W. Hu, David T. Ryan, Nicolas E. Grandel, Zacharie Maloney, George L. Britton, Maria A. Gonzalez Porras, Katerina Stojkova, Andrew Ligeralde, Hyeonwi Son, John Shannonhouse, Jacob T. Robinson, Aryeh Warmflash, Eric Brey, Yu Shin Kim, Amina A. Qutub

## Abstract

We introduce cytoNet, a cloud-based tool to characterize cell populations from microscopy images. cytoNet quantifies spatial topology and functional relationships in cell communities using principles of network science. Capturing multicellular dynamics through graph features, cytoNet also evaluates the effect of cell-cell interactions on individual cell phenotypes. We demonstrate cytoNet’s capabilities in four case studies: 1) characterizing the temporal dynamics of neural progenitor cell communities during neural differentiation, 2) identifying communities of pain-sensing neurons *in vivo*, 3) capturing the effect of cell community on endothelial cell morphology, and 4) investigating the effect of laminin α4 on perivascular niches in adipose tissue. The analytical framework introduced here can be used to study the dynamics of complex cell communities in a quantitative manner, leading to a deeper understanding of environmental effects on cellular behavior. The versatile, cloud-based format of cytoNet makes the image analysis framework accessible to researchers across domains.

**Availability and Implementation:** QutubLab.org/how | cytoNet contact: Brain Initiative Alliance Toolmaker cytoNet site: https://www.braininitiative.org/toolmakers/resources/cytonet/

**Author / Lay Summary:** cytoNet provides an online tool to rapidly characterize relationships between objects within images and video frames. To study complex tissue, cell and subcellular topologies, cytoNet integrates vision science with the mathematical technique of graph theory. This allows the method to simultaneously identify environmental effects on single cells and on network topology. cytoNet has versatile use across neuroscience, stem cell biology and regenerative medicine. cytoNet applications described in this study include: (1) characterizing how sensing pain alters neural circuit activity, (2) quantifying how vascular cells respond to neurotrophic stimuli overexpressed in the brain after injury or exercise, (3) delineating features of fat tissue that may confer resistance to obesity and (4) uncovering structure-function relationships of human stem cells as they transform into neurons.

## Introduction

Discoveries in biology increasing rely on images and their analysis (3). Advances in microscopy and accompanying image analysis software have enabled quantitative description of single-cell features including morphology, gene and protein expression at unprecedented levels of detail (4–7). There has also been a growing appreciation of spatial and density-dependent effects on cell phenotype. Various types of cell-cell interactions including juxtacrine and paracrine signaling are an integral part of biological processes that affect the behavior of individual cells. In response to this realization, many research groups have developed *in situ* profiling techniques to extract highly multiplexed single-cell data while preserving the spatial characteristics of biological samples (4, 8–12).

### Need for a user-friendly tool to test biological hypotheses that depend on spatial information

The increasing prevalence of spatially detailed imaging datasets has led to the proliferation of spatial analysis pipelines for biological research (**Table 1**). While these methods have enabled principled exploration of spatial hypotheses, the majority of the pipelines (with a few exceptions) have been developed for spatial molecular expression data obtained through methods such as mass cytometry, specialized high-resolution imaging, and/or scRNA-seq, with inherent idiosyncrasies. Others have focused on histology and samples obtained for medical applications. As a result, these techniques are not applicable to many standard imaging datasets obtained through routine biological experiments. Further, many pipelines require the user to be familiar with programming and involve the use of customized scripts. All of these limitations mean the most advanced spatial analysis platforms are not commonly employed by biologists. Instead, the spatial analysis platforms are largely used by a subset of labs heavily invested in computational analysis, by core users of specialized microscopy, or by imaging experts themselves. There remains a need for a generalizable, easy-to-use analysis method to test spatial hypotheses applicable to a wide variety of biological imaging data.

**Table 1.** Software tools for spatial analysis

| Software | Platform | Input | Output | Reference |
| --- | --- | --- | --- | --- |
| histoCAT | MATLAB,<br>standalone<br>program | Imaging mass<br>cytometry | User-guided cell neighborhood for<br>selected cells, enrichments/depletion of<br>cell-cell interactions based on comparison<br>to spatially randomized data | (50) |
| Pelkmans lab | Module<br>compatible with<br>CellProfiler | Cell cultures | Local cell density, population size, cell<br>islet edges | (34, 49, 51, 52) |
| Cell-graph | Standalone tool | H&E stained tissue<br>samples | Multiple graph metrics, e.g. clustering<br>coefficient, network diameter | (15) |
| PySpacell | Python | Cell cultures | Statistical tests of magnitude and scale of<br>spatial effects | (53) |
| SpatialDE | Python | Spatial transcriptomics<br>datasets | Statistical tests of genes with spatial<br>variation, spatial gene-clustering | (54) |
| trendsceek | R | Spatial transcriptomics<br>datasets | Statistical tests of genes with spatial<br>variation | (55) |
| cytoMAP | MATLAB | Histo-cytometry data | Multi-scale characterization of tissue<br>structure | (56) |
| MuSIC | Cytoscape | Immunofluorescence<br>and affinity<br>purification mass<br>spectrometry data | Intracellular protein positions and<br>distances | (57) |

### Need to capture time-dependency in structure-function relationships

In addition to spatial and morphological characteristics, time-dependent properties of cell function also define phenotype. The behavior of cell groups often includes coordinated responses of subgroups (such as in brain and heart tissue) that require intricate communication, and the role a cell plays in this communication is part of its phenotype. Live reporters and activity-based dyes can provide insight into this time-dependent cell communication. As an example, calcium imaging is a versatile technique to investigate the dynamics of cell signaling, particularly in neural and cardiac tissue. While there exist many automated tools for calcium signal analysis (**Table 2**), combined analysis of spatial and functional topology has the potential to reveal fundamental insight into the nature of structure-function coupling in biological systems.

**Table 2.**
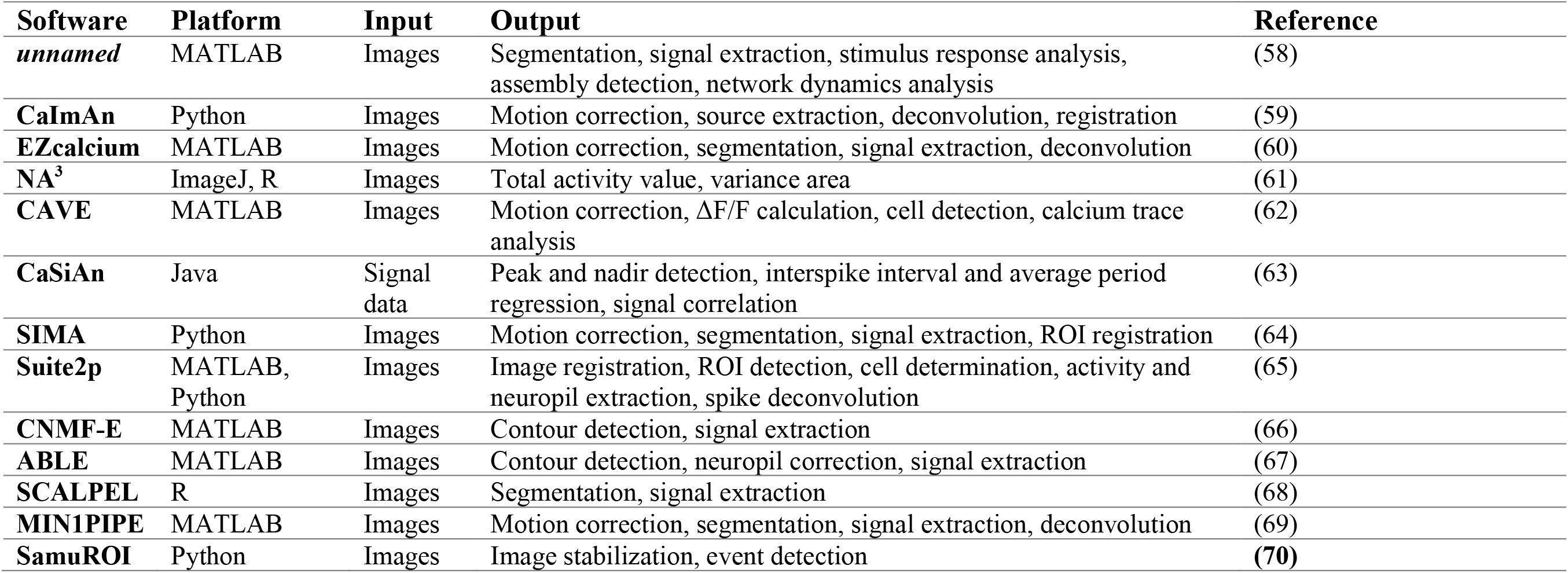
Software tools for calcium signal analysis.

| **Software** | **Platform** | **Input** | **Output** | **Reference** |
| --- | --- | --- | --- | --- |
| *unnamed* | MATLAB | Images | Segmentation, signal extraction, stimulus response analysis, assembly detection, network dynamics analysis | (58) |
| **CaImAn** | Python | Images | Motion correction, source extraction, deconvolution, registration | (59) |
| **EZcalcium** | MATLAB | Images | Motion correction, segmentation, signal extraction, deconvolution | (60) |
| **NA<sup>3</sup>** | ImageJ, R | Images | Total activity value, variance area | (61) |
| **CAVE** | MATLAB | Images | Motion correction, $\Delta F/F$ calculation, cell detection, calcium trace analysis | (62) |
| **CaSiAn** | Java | Signal data | Peak and nadir detection, interspike interval and average period regression, signal correlation | (63) |
| **SIMA** | Python | Images | Motion correction, segmentation, signal extraction, ROI registration | (64) |
| **Suite2p** | MATLAB, Python | Images | Image registration, ROI detection, cell determination, activity and neuropil extraction, spike deconvolution | (65) |
| **CNMF-E** | MATLAB | Images | Contour detection, signal extraction | (66) |
| **ABLE** | MATLAB | Images | Contour detection, neuropil correction, signal extraction | (67) |
| **SCALPEL** | R | Images | Segmentation, signal extraction | (68) |
| **MIN1PIPE** | MATLAB | Images | Motion correction, segmentation, signal extraction, deconvolution | (69) |
| **SamuROI** | Python | Images | Image stabilization, event detection | (70) |

### Network science framework

A single modeling framework to represent multiple descriptors of cell community is necessary to provide continuity across spatial and temporal scales. Network science offers this modeling framework. Network science seeks to understand complex systems by representing individual functional units of the system as nodes and their relationships as edges. This abstract representation is then used to describe, explain or predict the behavior of the system (13). Network models have been tremendously useful in studying complex biological systems, most prominently in neuroscience (13, 14). We posit that network models provide a flexible, intuitive method to model spatial and functional cell community relationships. Among existing image-based analyses that employ network science, the cell-graph technique (15) has been employed to great effect in analyzing structure-function relationships in fixed tissue sections. Our early work applying network analysis to fixed samples also enabled rapid classification of cell phenotypes (16, 17). However, the scope of network models in describing cell community structure and dynamics has yet to be fully explored.

Here we introduce cytoNet, a user-friendly method to analyze spatial and functional cell community structure from microscope images, using the formalism of network science (**Figure 1**). cytoNet is available as a web-based interface run on Amazon cloud. Users can choose to analyze image files from their desktops or online servers. Coupled with its ease-of-use, cytoNet’s versatility makes it accessible to researchers across domains. We originally designed the network modeling approach to study populations of developing neurons (2) and characterize how vascular cells respond to neurotrophic factors (16, 17). Here we extend the approach to case studies in a number of other biological systems. We partnered with labs from across research domains to illustrate applications of the cytoNet platform to stem cell biology, tissue engineering, and neuroscience in both *in vitro* and *in vivo* settings. The case studies demonstrate the broad utility of the network modeling approach in studying spatial and functional community structure in complex biological systems.

**Figure 1.**
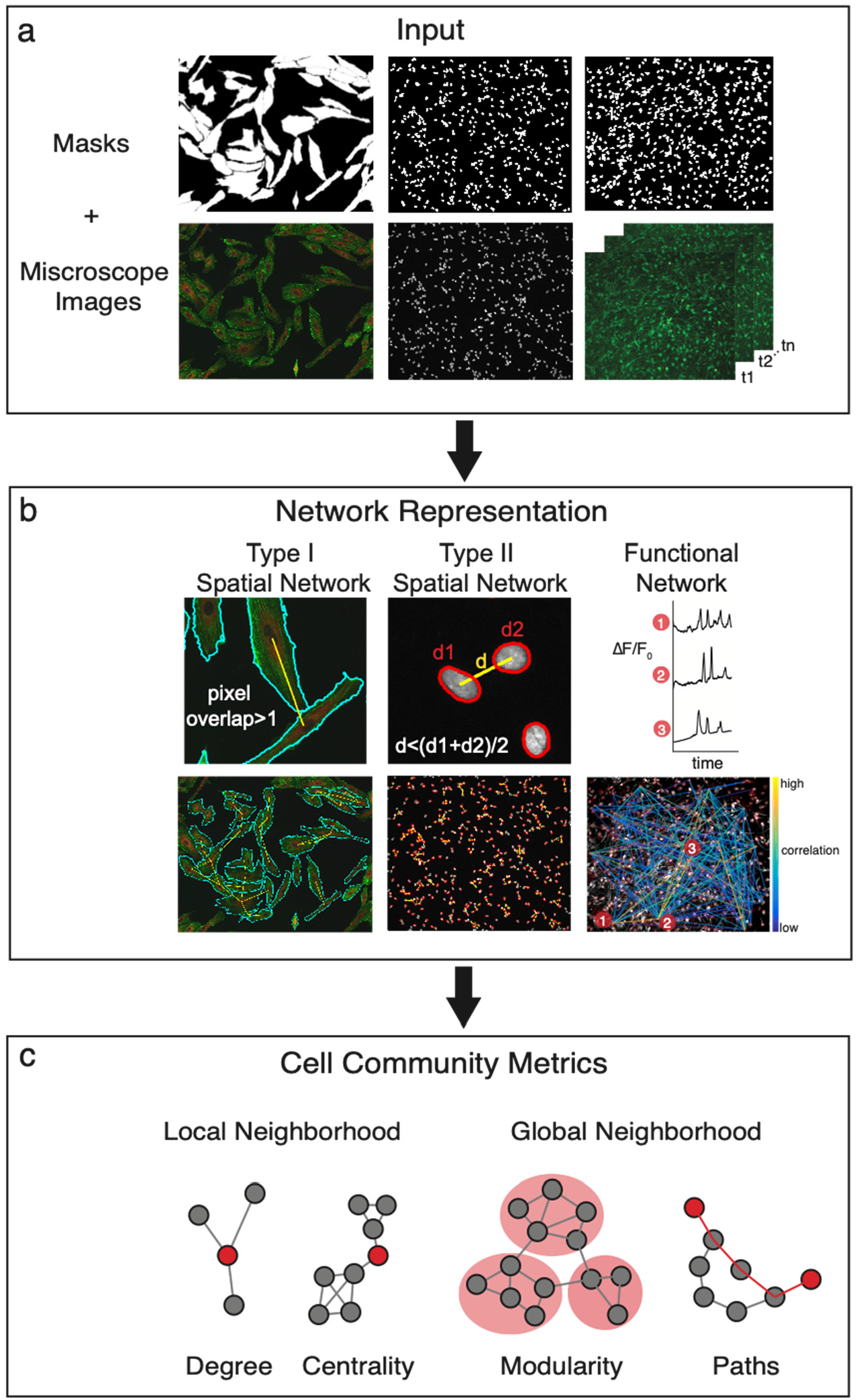
cytoNet workflow. (a) The cytoNet pipeline begins with masks and optionally microscope images, which can be static immunofluorescence images or calcium image sequences. (b) Spatial proximity is determined either by measuring shared pixels between cell pairs – type I networks, or by comparing the distance between cell centroids to a threshold distance – type II networks (right panel). Functional networks are estimated from correlations in calcium time series data. (c) Cell community descriptors provide information on local neighbrhood characteristics of individual cells, like degree and centrality measures, and global neighborhood characteristics like modularity and path lengths.

## Results

The cytoNet pipeline enabled us to investigate spatial and functional topology of cell communities in a variety of biological systems. Four case studies are described in the sections below.

### Case Study 1: Spatial and functional dynamics of neural progenitor cells (NPCs) during neural differentiation

We designed an *in vitro* model of neural differentiation to analyze the dynamics of spatial and functional topology during formation of neural circuits from neural progenitor cells (NPCs)^12^. NPCs are known to display structured intercellular communication prior to formation of synapses, which plays an important role in controlling self-renewal and differentiation (18–20). By leveraging the cytoNet method, we sought to capture the dynamic structure of NPC communities and the effect of such community structure on the phenotypes of individual cells.

In this case study, we describe data obtained using ReNCell VM human neural progenitor cells, in which spontaneous differentiation was triggered through withdrawal of growth factors, leading to rapid cell cycle exit and formation of dense neuronal networks in 5 days (2). We captured spontaneous calcium activity at days 1, 3, and 5 after withdrawal of growth factors. Following calcium imaging, cells were fixed, and nuclei were stained and reimaged. Nuclei images were then manually aligned by fiducial markers with their corresponding calcium images. The paired image sets allowed the creation of both functional and spatial graphs for the same communities of cells.

Spatial type II graphs (**Figure 2a**) showed a rise and fall in global network efficiency during neural differentiation (compared to randomized null models in which edges were rewired while preserving degree distribution; **Figure 2b)**. We hypothesize that these trends, independently confirmed in multiple NPC lines (2), reflect a transition from topologies favoring global to hierarchical information flow. We further explored this possibility through calcium imaging. Functional networks constructed from spontaneous calcium activity (**Figure 2c**) revealed network-wide signal correlations, with trends in spontaneous network activity mirroring spatial network parameters (**Figure 2d**). These results suggest that spatial topology predicts functional communication patterns in differentiating NPCs, with high spatial network efficiency at intermediate time points facilitating network-wide communication and low spatial network efficiency at early and late time points mirroring more clustered communication.

**Figure 2.**
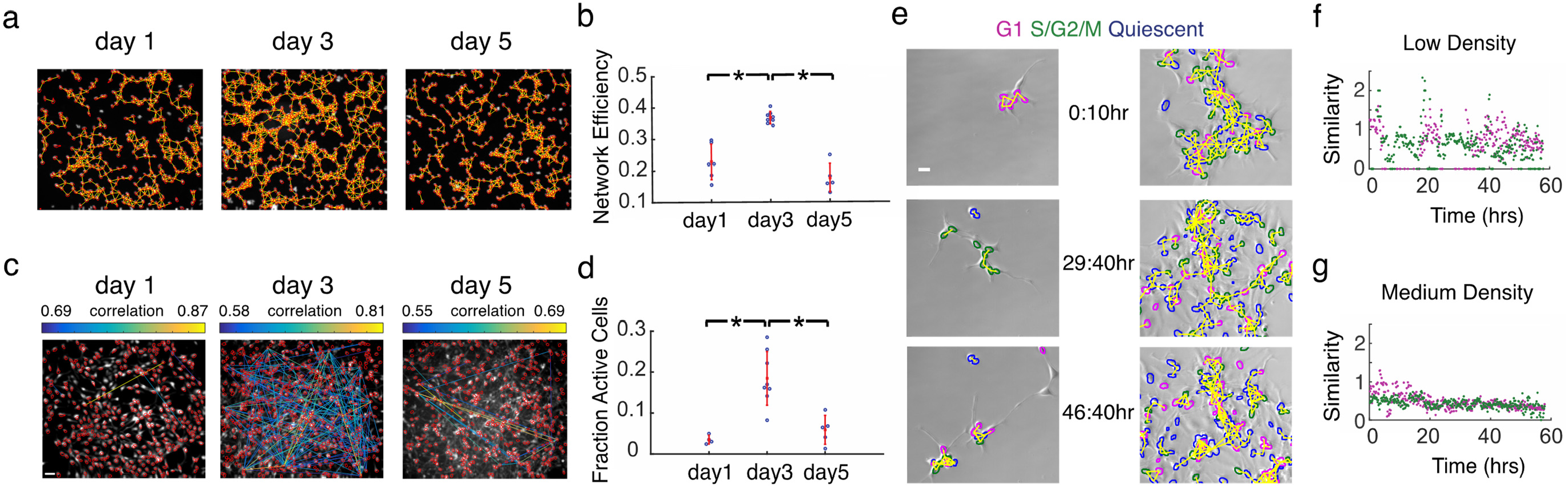
Dynamics of spatial and functional topology in developing neural progenitor cells (NPCs). (a) Spatial NPC networks at day 1, 3 and 5 of differentiation, overlaid on immunofluorescence images of nuclei stained with Hoescht dye; segmented cells are outlined in red, and spatial proximity edges are shown as yellow lines. (b) Network efficiency of spatial NPC networks peaks at day 3; red notches show mean and standard deviation; *p < 0.005 from two-sample t-test. (c) Functional networks obtained through calcium imaging with Fluo‐4 in developing NPC networks at days 1, 3 and 5. Correlations between calcium traces from individual cells are shown as a network plot overlaid on the maximum intensity image from calcium image sequences; scale bar = 50 μm for panels a and c. (d) Fraction of active cells in the network; *p < 0.005 from two-sample t-test. Active cells are defined as cells whose normalized fluorescence traces have three or more calcium transients. (e) Frames from time‐lapse movies of differentiating NPCs transfected with FUCCI cell cycle reporters. Borders of mCherry+ nuclei (G1) are outlined in magenta, Venus+ nuclei (S/G2/M) are outlined in green, and mCherry‐/Venus‐nuclei (quiescent) are outlined in blue, spatial edges are overlaid in yellow; scale bar = 50μm. (f) Neighborhood similarity score for low‐density culture across time. (g) Neighborhood similarity score across time for medium‐density culture. Figures 2a-d adapted from reference (2).

We next studied the role of cell-cell communication on cell cycle regulation of NPCs. Cell cycle regulation in NPCs is of interest as it has implications for the genetic basis of brain size in different species (21) and aberrant regulation can cause diseases like microcephaly (22). Studies in the ventricular zone of the embryonic mouse neocortex have shown that clusters of clonally-related neural progenitor cells go through the cell cycle together (23, 24). However, it is unclear whether this community effect is a ubiquitous feature of neural progenitor cells. To this end, we employed the cytoNet workflow to determine whether cell cycle synchronization is a feature of differentiating NPCs cultured in vitro.

For this part of the investigation, ReNCell VM human neural progenitor cells were stably transfected with the FUCCI cell cycle reporters (25) to generate Geminin-Venus/Cdt1-mCherry/H2B-Cerulean (FUCCI-ReN) cells. We captured time-lapse movies of FUCCI-ReN cells after withdrawing growth factors to induce differentiation and built network representations from nucleus images. Adjacency was determined by comparing centroid-centroid distance to a threshold (type II graphs).

In order to evaluate spatiotemporal synchronization in cell cycle, for each individual cell in a frame, we evaluated the average fraction of neighboring cells in a similar phase of the cell cycle (G1 phase – mCherry+ and S/G2/M phases – Venus+), normalized by total fraction of that cell type in the population. We called the average value of this fraction across all cells in an image the neighborhood similarity score, *N*_*S*_. Frames from time-lapse movies for low- and medium-density cultures are shown in **Figure 2e** (see also **Supplementary Videos 1-4**). We observed that groups of cells in the low-density culture moved through the cell cycle in unison, which was reflected in periodically high values of the neighborhood similarity score (**Figure 2f, Supplementary Video 1-2**). In contrast, the composition of cell clusters in the medium density culture was relatively heterogeneous, resulting in relatively low values of the neighborhood similarity score over time (**Figure 2g**, **Supplementary Video 3-4**). Neighboring cells in very low-density cultures are likely to be derived from the same clonal lineage, which explains the high level of synchronization in these cultures (23). This example highlights how cytoNet can be used to derive insight into the role of cell-cell interactions on dynamic cell behavior.

### Case Study 2: Dynamics of Coupled Functional & Spatial Analysis In Vivo

*In vivo* calcium analysis is an avenue for exploring and understanding the role that individual cells of the nervous system play in processing external stimuli including pain. Pain is mainly mediated by a subset of primary sensory neurons known as nociceptors in Dorsal Root Ganglia (DRG) and Trigeminal Ganglia (TG) (26). How DRG neurons function at a population level under physiological and pathological conditions is unknown. Imaging methods developed to record from hundreds to thousands of neurons simultaneously in the brains of live mice are helping elucidate this (27, 28). To investigate population characteristics of pain-sensing neurons, we used cytoNet to evaluate spatial and functional networks from calcium image sequences obtained in a mouse DRG model.

Calcium image sequences, along with single masks identifying individual cells, were inputs to cytoNet (see Methods for details on generation of masks) (**Figure 3**). Sensory stimulation experiments produced a single, major signal spike in each segmented cell (27). Measurement of the magnitude (ΔF/F0) of each spike is sensitive to the quality of segmentation; to mitigate this, we characterized each cell not by its spike magnitude, but by the time a cell took to reach its peak value from 20% of that value (ramp-up) and the time needed for the signal to return to 20% (ramp-down). Inspection of 44 segmented cells revealed 6 unique combinations of ramp-up times and ramp-down times (**Figure 3b**). Ramp-up times were either 5 or 10 seconds while ramp-down times varied between 5 and 35 seconds. This categorization of cells according to functional similarity was combined with the spatial graph of the segmented cells in order to identify spatial patterns of cells with similar behavior (**Figure 3a**). In addition, we note that although the vast majority of segmented cells reached their peak intensity at 20 seconds, a small group of cells along the left side of the tissue peaked at 25 seconds suggesting a right-to-left wave of response (**Figure 3**). This case study highlights the utility of cytoNet in analyzing spatial patterns of neural populations with unique functional signatures in an *in vivo* model.

**Figure 3.**
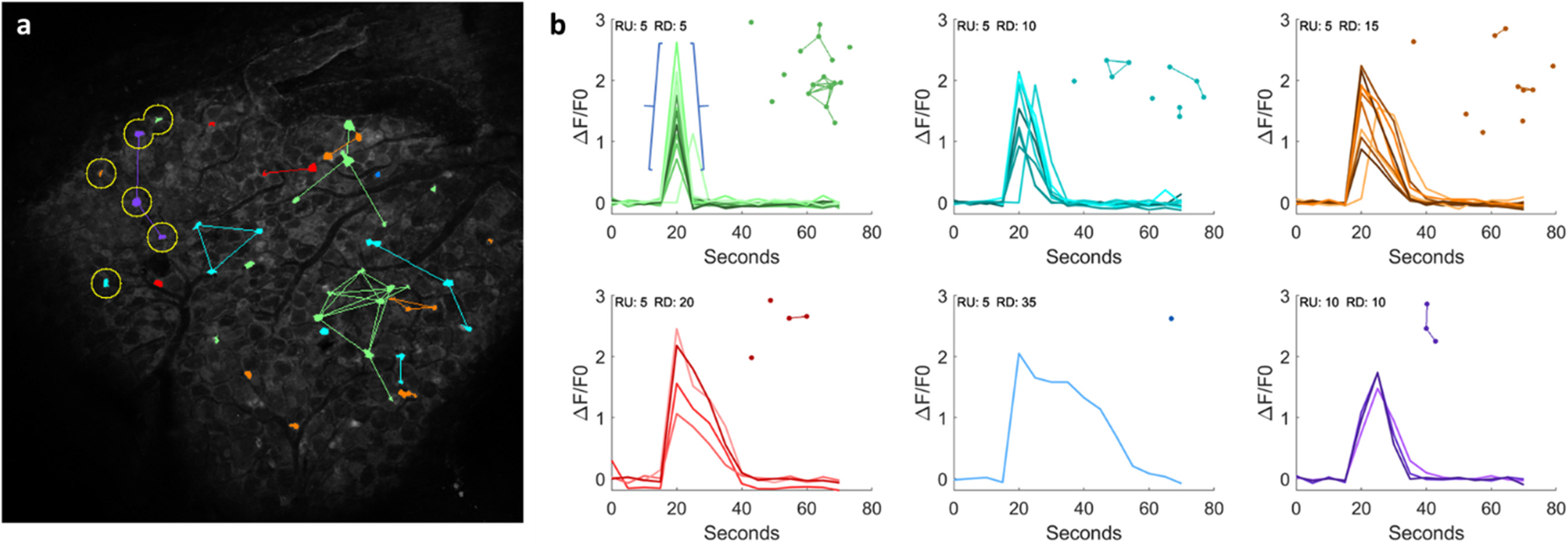
Dynamics of Coupled Functional & Spatial Analysis *In Vivo*. cytoNet captures relationships between spatial proximity of neurons and functional features of multicellular modules *in vivo*. (a) Cells classified according to the time required to first reach their maximum ΔF/F0 values from 20% of that value (ramp-up) and the time required to return to 20% (ramp-down). Edges connect similarly classified cells that are within 10 cell diameters of each other. All cells reached their peak values at 20 seconds except for those circled which reached their peak values at 25 seconds. (b) Calcium time series (ΔF/F0) plotted for 6 categories of cells with unique combinations of ramp-up and ramp-down times. The blue braces indicate a cell’s ramp-up and ramp-down. Each inset image is a spatial pattern of cells with the same ramp-up and ramp-down times. RU = ramp-up time; RD = ramp-down.

### Case Study 3: Disentangling the effect of cell community and growth factor stimulation on endothelial cell morphology

In a second application to studying human cells *in vitro*, we used cytoNet to evaluate the relative influence of local neighborhood density and growth factor perturbations on endothelial cell morphology. From a regenerative medicine perspective, studying the morphological response of endothelial cells to neurotrophic stimuli can help assess the cells’ potential angiogenic response following brain injuries that induce the secretion of neurotrophic factors, like ischemic stroke or transient hypoxia (29, 30). Common high-throughput angiogenic assays focus on migration and proliferation as the main cell processes defining angiogenesis, or the growth of new capillaries from existing ones (31). Distinct morphology and cytoskeletal organization of endothelial cells indicate the cell’s migratory or proliferative nature, and hence their angiogenic contribution within a sprouting capillary (32). Reproducibly quantifying the morphological response of endothelial cells to neurotrophic factors would enable more targeted approaches to enhancing brain angiogenesis.

We took an image-based approach to this problem, building a library of immunofluorescence images of human umbilical vein endothelial cells (HUVECs) stained for cytoskeletal structural proteins (actin, α-tubulin) and nuclei, in response to various combinations of vascular endothelial growth factor (VEGF) and brain-derived neurotrophic factor (BDNF) treatment. Cell morphology was annotated using 21 metrics described in our previous study (33) (**Supplementary Table 1**), which included cell shape metrics like circularity and elongation, and texture metrics for cytoskeletal stains such as actin polarity, smoothness etc. Network representations were designated based on shared cell pixels (type I graphs) and local network properties were described using the metrics in **Table 3.**

**Table 3.** Local neighborhood metrics calculated at the individual cell level ^1^Relevant only for type I graphs

| Graph Metrics | Symbol | Definition |
| --- | --- | --- |
| Degree | $k$ | Number of neighbors one link away from cell of interest |
| Average Neighbor Degree | $k_n$ | Average degree of all neighboring cells |
| Clustering Coefficient | $C$ | Number of edges in local neighborhood of a cell, divided by total possible connections |
| Local Efficiency | $E_l$ | Average shortest path length in local neighborhood |
| Node Closeness Centrality | $c_n$ | Sum of reciprocal distances in number of links to all other nodes |
| Node Betweenness Centrality | $w_n$ | Number of shortest paths that pass through a node |
| Shared Cell Border<sup>1</sup> | $S_b$ | Total number of pixels shared with neighbors |

<sup>1</sup> Relevant only for type I graphs

First, we quantified density-dependent effects on endothelial cell morphology in control cultures (without any growth factor perturbation). Our analysis showed correlations between cell morphological features and local network properties (**Supplementary Figure 3**). Some of these relationships were expected, for instance the positive correlation between shared cell border and cell size. Other relationships, such as the negative correlation between cell circularity and closeness centrality, capture intuitive notions of the influence of cell packing on morphology (**Figure 4a-c**). The closeness centrality of a cell (**Table 3**) describes its relative position in a colony cells in the middle of a colony will have higher centrality values than cells at the edge of a colony or isolated cells. The negative relationship between circularity and closeness centrality implies that isolated cells and cells located at the edge of colonies are more likely to have a circular morphology, while cells located at the center of colonies tend to be less circular (**Figure 4a-c**). Thus, our analysis revealed that local network properties have a quantifiable effect on cell morphology.

**Figure 4.**
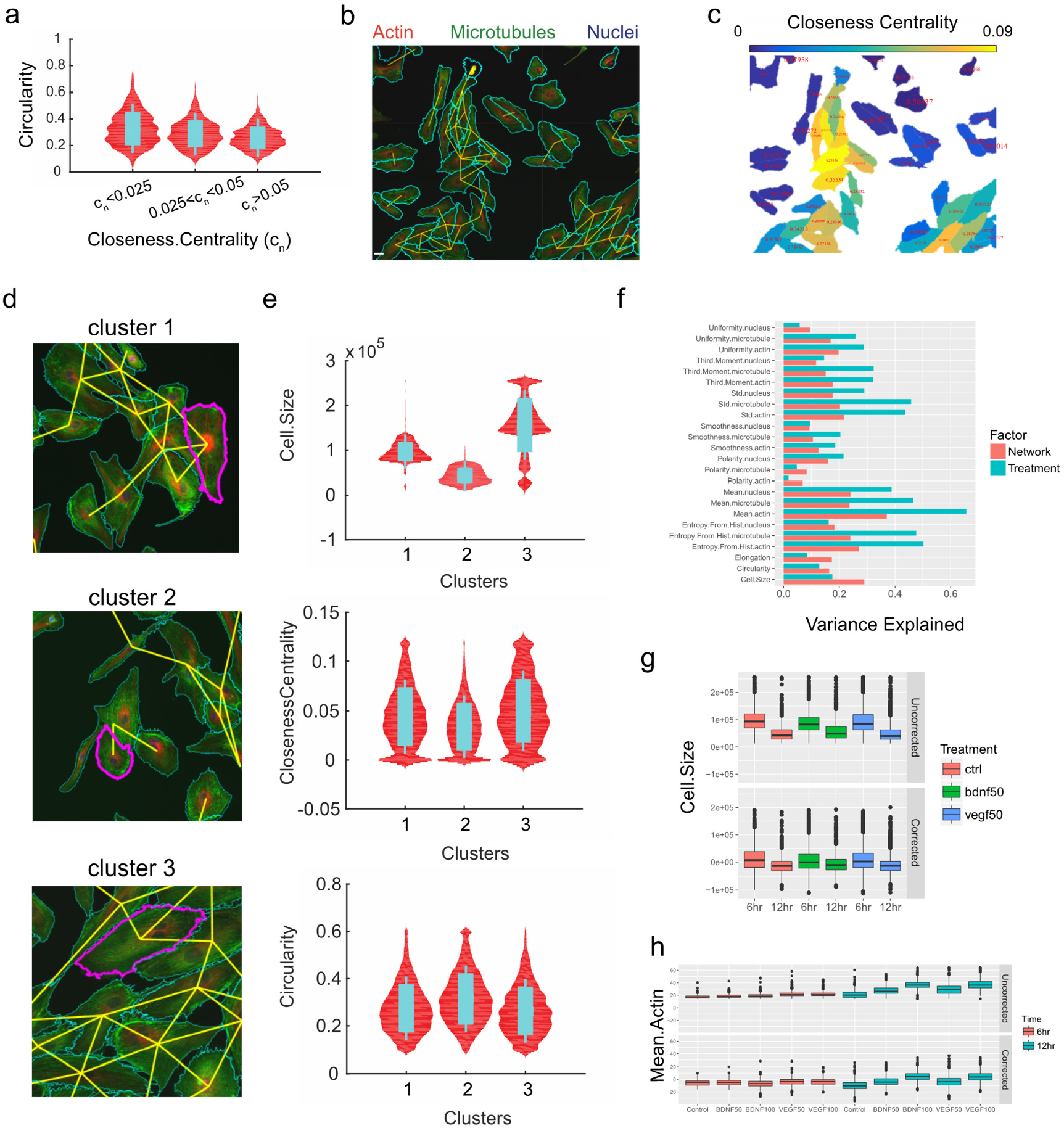
Influence of local neighborhood density on primary human endothelial cell (HUVEC) morphology. (a) Distribution of cell circularity values grouped under different levels of closeness centrality; sample size, n=786 cells (group 1; c_n_ < 0.025), 741 cells (group 2; 0.025 < c_n_ < 0.05) and 782 cells (group 3; c_n_ > 0.05); Cohen’s d effect size: groups (1, 2) = 0.34, groups (1, 3) = 0.62 **(b)** Sample immunofluorescence image with graph representation overlaid; scale bar = 50 μm. **(c)** Heatmap depicting closeness centrality of each cell, with circularity values overlaid in text. **(d)** Representative cells from cluster analysis, highlighted in magenta. **(e)** Cell size, closeness centrality and circularity distribution plots for each cluster. (**f)** Bar plot of variance explained by growth factor treatment and local network metrics. **(g)** Box plot of cell size as a function of growth factor treatment. **(h)** Box plot of mean actin intensity as a function of growth factor treatment. Legends and axes in (f-h) contain information on treatment (BDNF, VEGF), concentration (50ng/ml, 100ng/ml) and time of treatment (6 hours and 12 hours). Cohen’s d effect size for (f-h) is shown in **Supplementary Table 2**.

To determine dominant cell phenotypes, we performed cluster analysis on our dataset consisting of 25,068 cells. This analysis revealed 3 major categories of endothelial cells, with unique morphological and network signatures (**Figure 4d-e**). Cluster 1 comprised cells with migratory features, including low circularity and intermediate centrality indicative of their position at the edges of colonies. Cluster 2 contained small, circular cells with low centrality indicative of their isolation. Cells in cluster 3 showed proliferative features with large non-circular shapes, and high centrality indicating their positions in the center of colonies. Through this cluster-based phenotyping, we show how cytoNet can be used to infer the local environment and topological arrangement of distinct cell categories within a culture.

Next, we developed a workflow to analyze the effect of growth factor treatments on cell morphology, while correcting for the effect of local network properties. We did this to infer the independent effects of chemical perturbation and local cell crowding on cell morphology. First, we applied a quantile multidimensional binning approach (34, 35) to calculate the variance in morphology metrics that could be individually explained by all local network metrics and growth factor treatments (**Figure 4f**). We then calculated the values for each morphology metric after correcting for the effect of local network metrics (see Methods). The raw and network-corrected values for two metrics, cell size and mean actin intensity, are shown in **Figure 4g-h.** The influence of network properties can be clearly seen on cell size, where at 6 hours, large cell sizes are seen in the uncorrected but not corrected plots (**Figure 4g**). The effect of growth factor treatment can be clearly seen in network-corrected mean actin intensity (**Figure 4h**, **Supplementary Table 3**), where VEGF and BDNF treatment have dose-dependent effects on mean actin intensity independent of cell crowding effects. Thus, this case study demonstrates the utility of cytoNet in detecting the independent effects of local cell crowding and growth factor perturbations on cell morphology.

### Case Study 4: Spatial Analysis of the Pericapillary Niche in Adipose Tissue

In a second illustration of cytoNet’s utility to analyze intact tissue, we used cytoNet to characterize the pericapillary niche within adipose tissue. Specifically, we sought to understand the role of laminin α4, an extracellular matrix glycoprotein, in adipose tissue. Mice with a null mutation in the laminin α4 gene exhibit resistance to obesity and enhanced insulin sensitivity (36, 37). Understanding how the deletion of laminin α4 affects the spatial distribution of cells present in the adipose tissue can provide insight into the mechanisms underlying the functional change, and guide biomimetic models of the adipose perivascular niche (1, 38, 39). In this Case Study example, the confocal images of adipose tissue and capillaries were segmented by manual tracing on the computer, and provided as input to cytoNet. Because blood vessels have noncircular shapes, the distance between the centroids of vessels and other objects may not give a good sense of proximity. As an alternative graph-generation approach, cytoNet can compute the minimum distance between object perimeters in order to define graph edges. The resulting cell-to-cell perimeter distance table and cell area computations were used to determine differences between wild-type and knockout cells (**Figure 5**). The observed adipocytes stained with the BODIPY lipid dye tended to be smaller in knockout tissue compared to wild type (**Figure 5c**). This characterization is consistent with the observation that adipose in knockout mice is more similar to beige adipose tissue. In addition, we observed numerical differences in the “distance to capillary” metric for integrin α7 expressing cells between the laminin α4 knockout and wild-type mice models (**Figure 5f**), though for the limited sample size they were not statistically significant. Overall, these observations align with findings that the absence of laminin α4 leads to changes in stromal cell structure and distribution in pericapillary niches within adipose tissue (1). The resulting data can be used to guide studies into understanding the mechanisms underlying the effect of laminin α4 on adipose tissue function. Thus, this case study demonstrates the utility of cytoNet in detecting regional variations of cell structure within tissues and in addressing testable spatial hypotheses about tissue function.

**Figure 5.**
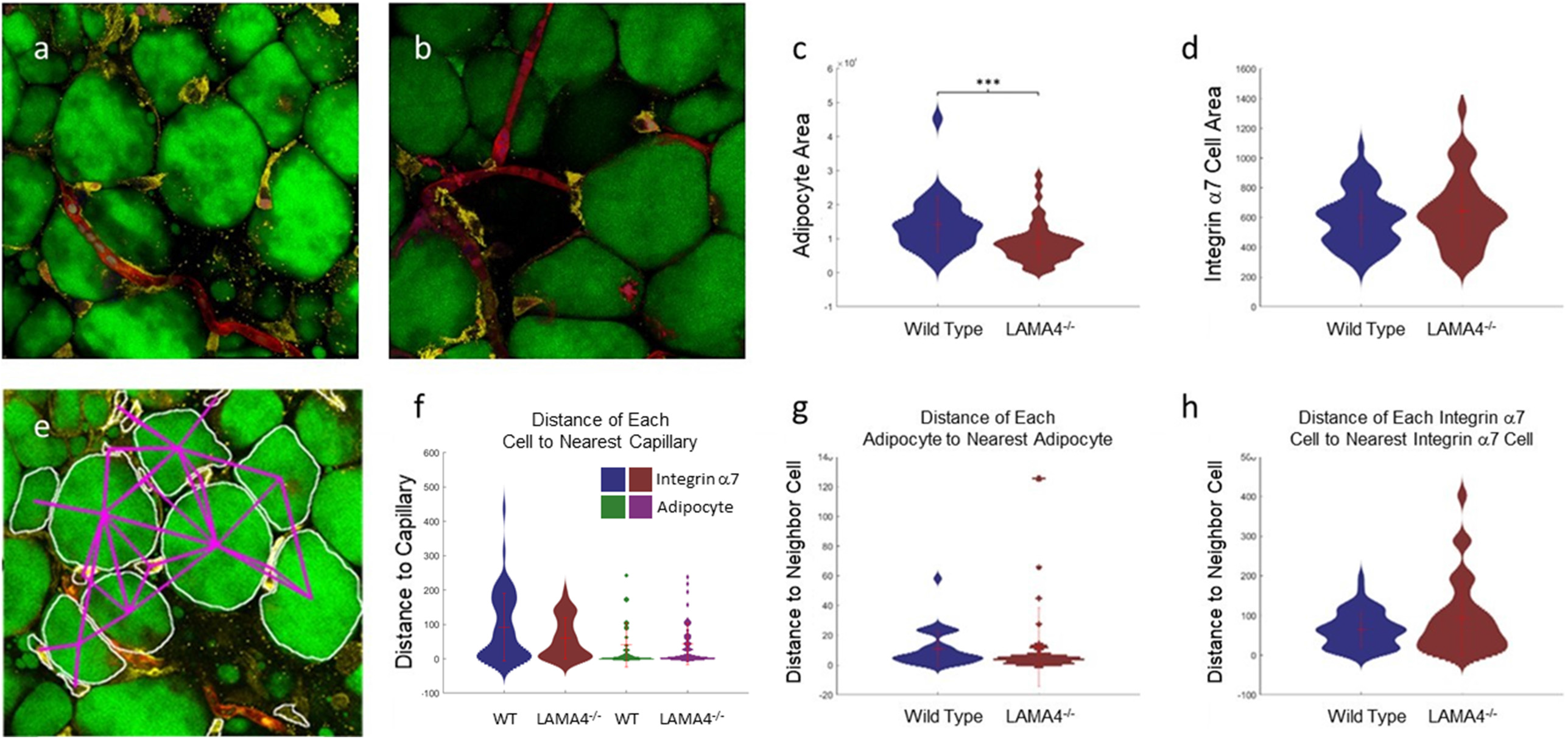
Spatial Analysis of the Pericapillary Niche in Adipose Tissue. Example confocal images of wild type (a) and knock out (b) adipose tissue and the corresponding output graph for the wild type image (e). Red = lectin (capillaries). Green = Bodipy (adipocytes). Yellow: integrin α7 positive cells. Violin plots of cell properties comparing wild-type and knockout (c, d, f-h). Distances are measured between the closest border pixels of pairs of objects. Figure 5f is adapted from reference (1). Error bars are mean +/-standard deviation. p-values were computed using the Wilcoxon rank sum test (***: p ≤ 0.001).

## Discussion

Advances in *in situ* profiling techniques have led to the generation of highly multiplexed imaging datasets describing tissue archicture in great spatial detail (4, 8–12). Spatially detailed imaging datasets have led to a proliferation of computational pipelines designed to test spatially driven biological hypotheses (**Table 1**). However, many of these analysis pipelines are designed specifically for spatial molecular expression data and are not generalizable to data obtained from other microscopy techniques. Further, due to their reliance on specialized scripts, many pipelines are not readily accessible to biological researchers without programming background.

Here we present cytoNet, a user-friendly pipeline for investigation of spatial hypotheses in cell- and tissue-based biological experiments. cytoNet is available through an intuitive web interface, eliminating the need to download and install software. Source code is also provided as MATLAB scripts for more advanced users. Pre-segmented masks provided as input to cytoNet are used to build network representations of spatial topography. Accompanying fluorescence or confocal images are used to extract single-cell features and functional relationships. Lastly, network descriptors are combined with single-cell features to explore cell community effects on cell phenotypes.

We demonstrate the utility of cytoNet through four case studies. As shown in detail in our previous study (2), we harness an *in vitro* model of neuronal network formation from neural progenitor cells (NPCs) to demonstrate a rise and fall in network efficiency during neural differentiation. Accompanying functional network analysis through calcium imaging shows that these trends in community structure likely reflect a transition from global to hierarchical communication during the formation of neural circuits. We further use local neighborhood measures to explore the effect of cell community on cell cycle regulation, showing a density-dependent effect on cell cycle synchronization.

Our second case study showed cytoNet’s capability for analyzing time-varying functional image sets. In this case, we characterized spatiotemporal calcium signaling recorded from intact brain tissue. Networks can be constructed based on the similarity of temporal behaviors of cells. The combination of the functional networks and spatial networks reveals local groups of cells with similar behaviors and assists in the development and testing of hypotheses of functional subsystems in neuronal tissue.

We also explored the differential effects of cell density and growth factor stimulation on human endothelial cells using cytoNet. By applying unsupervised clustering approaches on a suite of cytoNet-generated metrics describing cell morphology and local neighborhood, we show the presence of three cell phenotypes. These phenotypes reflect different cytoskeletal states and multicellular interactions indicative of collective behaviors like migration and proliferation. Further, we leverage a quantile multidimensional binning approach to investigate the differential effects of cell density and growth factor perturbations on cell morphology. This workflow can be used to comprehensively characterize the response of cells to chemical perturbations and aid in drug discovery. Case Study 4 illustrated another translational application of cytoNet: this time to study the effect of an extracellular matrix protein on the phenotype of adipose cells within perivascular niches.

Notably, two of the case studies were applied in vitro to human cells, and two were applied to *in vivo* image sets. Case Study 1 and 2 capitalized on cytoNet’s ability to integrate functional and structural graphs across time in a single mathematical framework. The other two cases illustrated the how cytoNet can be applied to optimize cell phenotyping (Case Study 3 and 4). All of the cases show how cytoNet can help guide hypotheses, inform biomimetic models or tailor therapeutic interventions that reflect a cell’s microenvironment.

The network model utilized by cytoNet is a versatile modeling framework that can be used to incorporate many hypotheses on cell-cell interactions and their role in cellular behavior. In future iterations, this framework can be expanded to incorporate non-binary interactions through weighted networks, shift the focus from individual nodes to motifs through simplicial complexes, and include dynamic reconfiguration of networks over time through multilayer networks. Further, once graphs have been defined, graph theory affords a rich array of metrics that can be used to probe network structure, only some of which were studied here. These include a variety of null graph models that can be used to test specific spatial hypotheses.

In summary, the cytoNet method provides a user-friendly spatial analysis software, leveraging network science to model spatial topography and functional relationships in cell communities. This framework can be used to quantify the structure of multi-cellular communities and to investigate the effect of cell-cell interactions on individual cell phenotypes.

## Methods

### Software

cytoNet is available as a web-based interface at https://www.QutubLab.org/how and associated scripts are available at https://github.com/arunsm/cytoNet-master.git. An overview of cytoNet as a resource for the Brain Initiative Alliance community is provided here, along with video tutorials: https://www.braininitiative.org/toolmakers/resources/cytonet/

See **Supplementary Methods 1** for instructions on using cytoNet.

### cytoNet image analysis pipeline

The cytoNet pipeline begins with masks and accompanying microscope images. The microscope images may be any color or gray-scale based microscopy images (e.g., immunofluorescence, confocal) or a sequence of calcium images (**Figure 1a**). The provided mask is used to extract features of cells and to construct spatial and functional graphs (**Figure 1b**). Spatial graphs are created by having nodes represent mask objects and edges determined by object distance. Edges can be found by one of two methods for spatial graphs: by evaluating the distance between cell boundaries (type I graphs), or by evaluating the proximity of cells in relation to a threshold distance (type II graphs) (**Figure 1b**). The type I graphs are useful when detailed information of cell boundaries and morphology is available, such as in the case of membrane stains or cells stained for certain cytoskeletal proteins. The type II graphs work well with images of cell nuclei, where detection of exact cell boundaries is not possible. In both approaches, cells deemed adjacent to each other are connected through edges, resulting in a network representation. If calcium imaging sequences are provided as input, a functional graph is created based on correlations among calcium time series of different mask objects (**Figure 1b**).

### Image Segmentation

Image segmentation – the identification of salient foreground objects such as cells – is often the first step in image analysis. The cytoNet pipeline works with pre-segmented masks of images and accompanying microscope images. For users who do not have mask files, cytoNet includes basic image segmentation algorithms including thresholding and watershed operations to generate these masks. The segmentation algorithms included in cytoNet can be parameterized to work well for images with clear delineation of nuclei and cell borders, like the endothelial cell examples provided on the cytoNet website. The cytoNet code also provides frequency detection of cells, where a change in a functional marker (e.g., Ca^2+^ or FUCCI) delinates cell location. For object detection in most other image sets, we point the user to programs that focus on cell segmentation (40–42). Multiple research teams have made significant inroads into designing generalizable image segmentation algorithms, among them classic thresholding and watershed approaches (43), pixel-based classifiers (40) and more recently deep learning approaches (4, 41, 42). These programs generate masks as output. Users may wish to implement them prior to analyzing community structure through cytoNet. Image segmentation and graph creation are handled separately by cytoNet, enabling flexibility for the user.

### Generation of spatial networks

Type I graphs are generated as follows. Mask boundaries are expanded by 2 pixels and overlap of expanded masks is used to assign edges and build an adjacency matrix. Cells touching the image border are included in calculations of local network properties (**Table 3**) for cells not touching the boundary but are excluded for the construction of the adjacency matrix. Type II graphs are generated as follows: for each pair of objects (nuclei), a threshold distance for proximity is defined as the average of the two object diameters, multiplied by a scaling factor (S). If the Euclidean distance between the object centroids is lower than the threshold distance computed, the pair of objects is connected with an edge. We chose a default scaling factor S = 2 for all our analyses, through visual inspection of cell adjacency.

### Generation of functional networks

Functional networks are created using the method described by Smedler et al, (44) where cross-covariance between signals is used to assign functional connections between pairs of cells (**Figure 1b**). A randomized dataset is generated by shuffling each signal in the original dataset at a random time point. The 99^th^ percentile of cross-covariance values for the randomized dataset is used as a threshold for determining significant correlations.

### Network Metric Computation

For both spatial and functional graphs, connectivity is denoted mathematically using an adjacency matrix, *A*, where *A*_*i*,*j*_ = 1 if there exists an edge between cells *i* and *j*, and 0 otherwise. This concise representation of hypothesized interactions among cells can be used to generate multiple descriptors at a local level for individual nodes and at a global level for the entire graph (**Figure 1c**). Extracted metrics are used to visualize and analyze local neighborhood effects on individual cell phenotypes (**Table 3**), as well as global cell community characteristics (**Table 4**). Examples of local metrics are number of connections (degree) or notions of centrality, such as ability to act as a bridge between different cell communities (betweenness centrality). Examples of global metrics include measures of modularity such as the number of connected components, and measures of information flow such as path length. All the network metrics described in **Table 3** and **Table 4** were computed using custom-written code, building upon routines provided in (45).

**Table 4.**
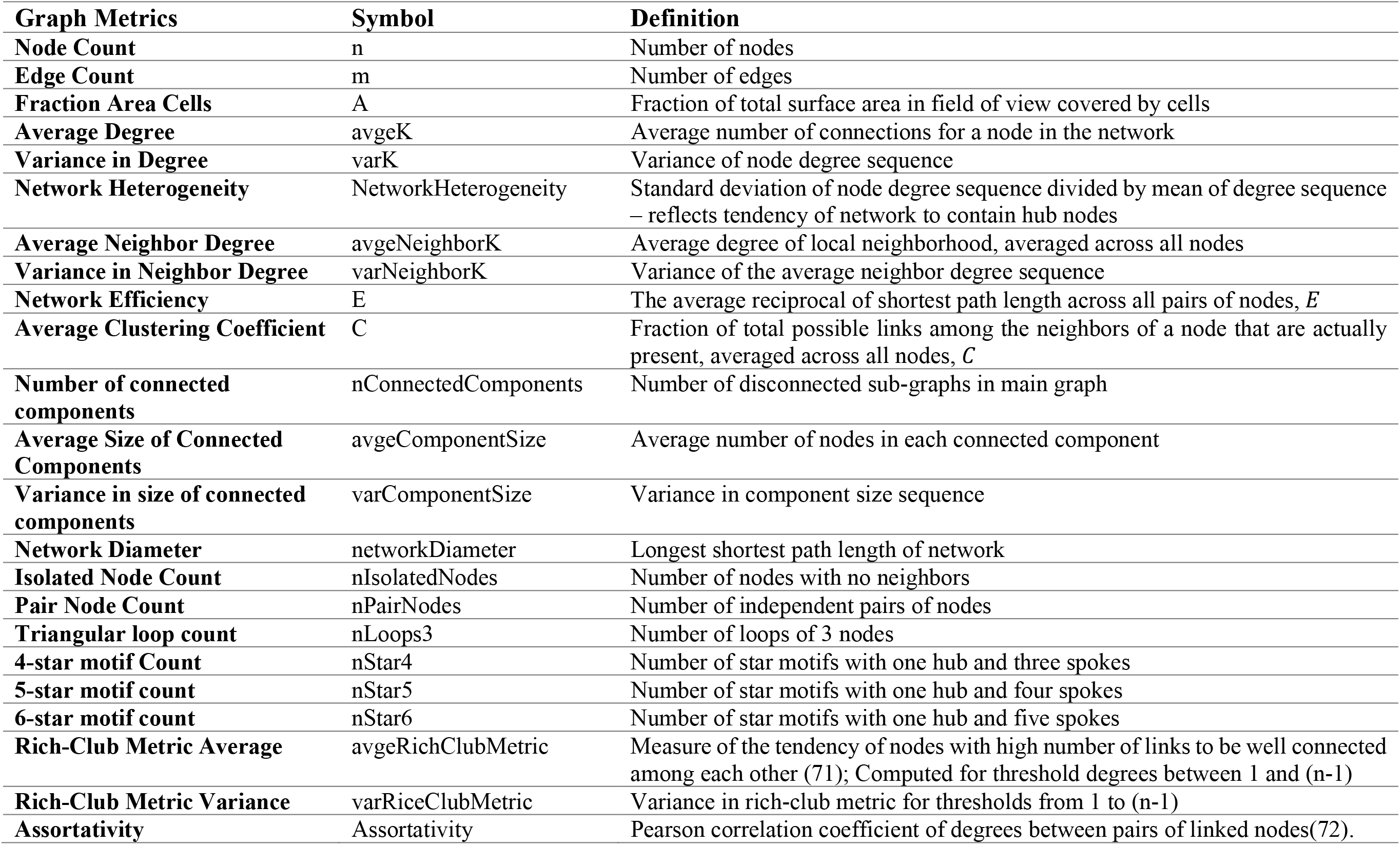
Global graph metrics and their normalization to account for network size. n = number of nodes, m = number of edges.

### Cell Culture

Human umbilical vein endothelial cells (HUVEC) were obtained from Lonza and cultured in EBM-2 medium (Lonza) supplemented with penicillin-streptomycin (Fisher Scientific) and EGM-2 SingleQuot bullet kit (Lonza). For imaging experiments, cells were cultured for different periods (6, 12 or 24 hours) in different combinations of vascular endothelial growth factor (VEGF, human recombinant; Millipore) and brain-derived neurotrophic factor (BDNF, human recombinant, Sigma-Aldrich). Concentrations used were in the range 50-100 ng/ml. Controls were the same culture period without growth factor treatments.

Immortalized human neural progenitor cells derived from the ventral midbrain (ReNCell VM) were obtained from Millipore. Cells were expanded on laminin-coated tissue culture flasks, in media containing DMEM/F12 supplemented with B27 (both Life Technologies), 2 μg/ml Heparin (STEMCELL Technologies), 20 ng/ml bFGF (Millipore), 20 ng/ml EGF (Sigma) and penicillin/streptomycin. For differentiation experiments, cells were cultured in medium lacking bFGF and EGF.

### Dorsal Root Ganglion Mouse Model

Dorsal laminectomies were performed on anesthetized mice exposing the dorsal root ganglia in the spinal L5 region. The spinal columns were stabilized under a laser-scanning confocal microscope. Stimuli were applied to the hind paw in one of four ways: 1) pressure (rodent pincher analgesia meter), 2) gentle mechanical stroke (brush or von Frey filament), 3) thermal stimuli (immersion in hot or cold water), 4) chemical stimuli (KCl, capsaicin, or TRPV1 agonist applied subcutaneously). Calcium image sequences were acquired at depths of up to 100 μm at 1-3 Hz at intervals of 4-6 seconds.

### Laminin α4 Knockout Mouse Model

Subcutaneous fat was separately collected from laminin α4 knock out mice and wild-type mice. The samples were processed and incubated with integrin α7 antibody (1:100, Novus Biologics NBP1-86118) and Griffonia simplicifolia isolectin conjugated with Rhodamine (labels endothelial cells/blood vessels) followed by incubation with a second antibody (Alexa Fluor 647 Donkey Anti-Rabbit IgG, Abcam ab150075) and BODIPY to stain lipid. Images were collected by a Leica TCS SP8 Confocal Microscope.

### NPC calcium image acquisition and processing

ReNCell VM neural progenitor cells were plated on LabTek chambered cover glasses for calcium imaging experiments. Cells were loaded with culture medium containing 3 μM of the fluorescent calcium indicator Fluo-4/AM (Life Technologies) and Pluronic F-127 (0.2% w/v, Life Technologies) for 30 min at 37°C. Imaging of spontaneous calcium activity was performed at 37°C using a 20X objective lens (N.A. = 0.75), with 488 nm excitation provided through a SOLA SE Light Engine (Lumencor). 16-bit fluorescence images were acquired at a sampling frequency of 1 Hz for a total duration of 15 min, using a Zyla 5.5 sCMOS camera (Andor). Following calcium imaging, samples were fixed, and nuclei were stained using DAPI. By navigating to the locations where calcium imaging was performed, manual co-registration was done to obtain immunofluorescence images for the same fields of view.

Regions of interest (ROIs) were obtained by segmenting nucleus images using a local thresholding approach followed by the watershed algorithm. Undersegmented objects were algorithmically removed by discarding the top two percentile of object sizes obtained after segmentation. Next, a time-varying fluorescence trace was calculated for each ROI. For each frame in the calcium fluorescence image stack, background (average pixel intensity of non-ROI regions in the image) was subtracted. Average fluorescence intensity for each ROI (*F*) was obtained by averaging pixel intensity values within the ROI for each time point. Baseline fluorescence (*F*_*0*_) for each ROI was calculated as the minimum intensity value in a window 90s before and after each time point. The normalized fluorescence trace for the ROI was then calculated as *F* − *F*_0_/*F*_0_. Cells with low activity were filtered out by discarding traces with less than three peaks and traces whose signal-to-noise ratio was lower than 1. Quality of the remaining traces was confirmed by manual inspection. This was done to avoid false positives in the cross-correlation analysis.

### Generation of FUCCI Reporter Neural Progenitor Cell Lines

Stable reporter cell lines (FUCCI-ReN) were generated by sequentially nucleofecting ReNcell VM neural progenitor cells with an ePiggyBac (46) construct encoding mCherry-Cdt, Venus-Geminin, or Cerulean-H2B. Each construct introduced to the cells was driven by a CAG promoter containing a blasticidin (ePB-B-CAG-mCherry-Cdt1), puromycin (ePB-P-Venus-Geminin), or neomycin (ePB-N-Cerulean-H2B) resistance gene. Following each round of nucleofection, cells were cultured in the presence of appropriate antibiotics (2 μg/ml blasticidin, 0.1 μg/ml puromycin and 100 μg/ml neomycin).

### Acquisition and processing of FUCCI-ReN time lapse videos

FUCCI-ReN cells were plated at different densities on chambered cover glasses (Fisher Scientific) coated with laminin. Cells were imaged after switching to differentiation medium containing phenol red-free DMEM/F12. Time-lapse imaging was performed using a Nikon Ti-E microscope equipped with a motorized stage, a cage incubator for environmental control (Okolab), a 20X objective lens (N.A. = 0.75), SOLA SE Light Engine for LED-based fluorescence excitation (Lumencor), appropriate filters for visualizing mCherry, Venus and Cerulean fluorescent proteins and a Zyla 5.5 sCMOS camera (ANDOR). 16-bit composite fluorescence images were acquired at 10-minute intervals for a total duration of 57.5 hours.

Grayscale images for each channel (H2B-Cerulean, Geminin-Venus and Cdt1-mCherry) were binarized using locally adaptive thresholding. Seeds for the watershed transform were generated using the regional minima from the distance transform of the grayscale images. Next, the watershed algorithm was applied to detect boundaries between overlapping cell nuclei. Finally, information from different channels were used to correct undersegmented nuclei (**Supplementary Figure 2**).

### Acquisition and processing of HUVEC immunocytochemistry images

For imaging experiments, HUVECs were cultured on glass dishes coated with fibronectin (Sigma-Aldrich). After appropriate growth factor treatments, cultures were fixed with 4% paraformaldehyde, free aldehyde groups were quenched using 1 mg/mL sodium borohydride, and membranes were permeabilized with 0.2% Triton-X-100 solution in PBS. Actin fibers were visualized using an Alexa Fluor 488-phalloidin antibody (1:40, Molecular Probes) and microtubules were visualized using a mouse monoclonal anti-α-Tubulin antibody (1:250, Sigma-Aldrich) followed by a goat anti-mouse Alexa Fluor 647 secondary antibody. Nuclei were stained using Hoescht (Molecular Probes). 16-bit composite immunofluorescence images were acquired through a 20X objective (N.A. = 0.75) on a Nikon Ti-E epifluorescence microscope. Physical pixel size was 0.32 μm.

Fluorescence images were processed as described previously (47) (**Supplementary Figure 1**). Briefly, the following steps were used.

1. Contrast was enhanced using histogram equalization.
2. Images were smoothed using a 2D Gaussian lowpass filter.
3. Initial binarization was performed using Otsu’s method.
4. The binary image was dilated to fill in individual cell areas.
5. All objects <1% of the total image area were removed. This was called the final binary image.
6. A binary representation of the nuclear and microtubule image layers was generated using a high input threshold value. This was called the marker image.
7. Another binary image was created with values of 0 where either the final binary image (step 5) or the marker image (step 6) had a value of 1.
8. Watershed markers were generated by imposing the minimum of the complement of images obtained in steps 2 and 7. This image had black markers contained within cells to serve as basins for flooding, while cell areas themselves were represented by lighter pixels that served as the rising contours of the basins.
9. The watershed algorithm was implemented using Matlab’s built-in function to generate cell boundaries.
10. Masks generated in step 9 were refined by using composite images of microtubules and actin as the marker image (step 6).

In order to automate the threshold generation, the area of cell masks obtained from segmentation were compared to those obtained through thresholding with a high threshold. The entire process was then iterated until an acceptable area ratio was achieved.

### Processing of In Vivo Calcium Image Sequences

Calcium image sequences from dorsal root ganglion models were processed as follows. To generate a mask, the calcium image sequence was first decomposed into individual grayscale frames. Next, for each pixel location, the maximum and minimum intensities were found across all frames. The differences between the maximum and minimum intensities were stored in an array (of delta values) and normalized. An initial segmentation of the delta values was done by thresholding using Otsu’s method, resulting in an initial binary mask. The initial mask was refined by computing a new threshold by applying Otsu’s method to only those delta values that were identified as foreground objects in the initial segmentation. The resulting binary image underwent a morphological closing with a disk of radius 3, and objects of fewer than 10 pixels were removed to generate the final mask.

To generate functional networks, edges were placed between two cells whenever: a) the two cells had the same ramp-up and ramp-down times, and b) the Euclidean distance between the centroids of the two cells was less than or equal to 10 times the mean of the diameter of each of the two cells.

### Cluster Analysis

We performed cluster analysis on the HUVEC imaging dataset using Shrinkage Clustering (48), a two-in-one clustering and cluster optimization algorithm based on matrix factorization that simultaneously finds the optimal number of clusters while partitioning the data. Cells whose features had the smallest sum of squares distance to the median values for each cluster were identified as representative cells for each cluster.

### Correction of Morphology Metrics for Effects of Local Network Properties and Treatment Conditions

We performed quantile multidimensional binning (49) of cells for all 7 network metrics (5 bins per metric). The mean of each morphology metric was calculated for each multidimensional bin, and this mean was subtracted from the raw measurements to generate the network-corrected measurements for each cell. Treatment-corrected measurements were generated similarly by calculating the mean of each morphology metric under each treatment condition and then subtracting it from the raw measurements.

### Variance Explained by Local Network Properties and Treatment Conditions

The variance explained by each factor was calculated using the following formula (35)

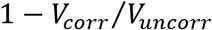

*V*_*corr*_ is the variance of the corrected measurements, and *V*_*uncorr*_ is the variance of the uncorrected measurements.

## Acknowledgements

We thank Dr. David Noren, Dr. André Schultz and Dr. Tien Tang for helpful discussions and comments on the manuscript, and Amada Abrego, Dr. Becky Zaunbrecher and Grace Ching for technical assistance. We also thank the Brain Alliance Initiative for curating cytoNet as a resource for the research community, the Keystone Symposia for highlighting cytoNet as a virtual SciTalk resource. This work was supported by NSF Career Grant 1150645 to AAQ, NSF Neural and Cognitive Systems grant 1533708 to AAQ and JTR, NSF award 1553228 to AW, CPRIT award RR140073 to AW, and National Institutes of Health Grant RO1DE026677 and UT Rising STAR Award to YSK. ASM was supported through NSF IGERT training grant 1250104.

## Author Contributions

ASM, DTR, EB, YSK and AAQ designed the experiments. ASM, GLB, DTR, MGP, KS, HS and JS performed the experiments. BLL, ASM, CWH, NEG, ZM and AAQ analyzed the data. BLL, AL and AAQ designed and implemented the cytoNet website. All authors contributed to writing the manuscript. AAQ, AW, EB and YSK supervised the work.

## References

1. Gonzalez-Porras M, Stojkova K, Vaicik MK, Pellowe A, Goddi A, Carmona A, et al. Integrins and extracellular matrix proteins module adipocyte thermogenic capacity. Scientific Reports. 2021;in press.

2. Mahadevan AS, Grandel NE, Robinson JT, Francis KR, Qutub AA. Living Neural Networks: Dynamic Network Analysis of Developing Neural Progenitor Cells. bioRxiv. 2021.

3. Meijering E, Carpenter AE, Peng H, Hamprecht FA, Olivo-Marin J-C. Imagining the future of bioimage analysis. Nature Biotechnology. 2016;34(12):1250–5.

4. Mund A, Coscia F, Hollandi R, Kovács F, Kriston A, Brunner A-D, et al. AI-driven Deep Visual Proteomics defines cell identity and heterogeneity. bioRxiv. 2021:2021.01.25.427969.

5. Bray M-A, Singh S, Han H, Davis CT, Borgeson B, Hartland C, et al. Cell Painting, a high-content image-based assay for morphological profiling using multiplexed fluorescent dyes. Nature Protocols. 2016;11(9):1757–74.

6. McQuin C, Goodman A, Chernyshev V, Kamentsky L, Cimini A, Karhohs KW, et al. CellProfiler 3.0: Next-generation image processing for biology. PLoS Biology. 2018:1–17.

7. Kuthuru S, Szafran AT, Stossi F, Mancini MA, Rao A. Leveraging Image-Derived Phenotypic Measurements for Drug-Target Interaction Predictions. Cancer Inform. 2019;18:1176935119856595.

8. Schürch CM, Bhate SS, Barlow GL, Phillips DJ, Noti L, Zlobec I, et al. Coordinated Cellular Neighborhoods Orchestrate Antitumoral Immunity at the Colorectal Cancer Invasive Front. Cell. 2020;182(5):1341–59.e19.

9. Vickovic S, Eraslan G, Salmén F, Klughammer J, Stenbeck L, Schapiro D, et al. High-definition spatial transcriptomics for in situ tissue profiling. Nature Methods. 2019;16(10):987–90.

10. Gut G, Herrmann MD, Pelkmans L. Multiplexed protein maps link subcellular organization to cellular states. Science. 2018;361(6401).

11. Eng CHL, Lawson M, Zhu Q, Dries R, Koulena N, Takei Y, et al. Transcriptome-scale super-resolved imaging in tissues by RNA seqFISH+. Nature. 2019;568(7751):235–9.

12. Thul PJ, Åkesson L, Wiking M, Mahdessian D, Geladaki A, Ait Blal H, et al. A subcellular map of the human proteome. Science. 2017;356(6340).

13. Bassett DS, Zurn P, Gold JI. On the nature and use of models in network neuroscience. Nature Reviews Neuroscience. 2018;19(9):566–78.

14. Bassett DS, Sporns O. Network neuroscience. Nature Neuroscience. 2017;20(3):353–64.

15. Yener B, Bülent. Cell-graphs. Communications of the ACM. 2016;60(1):74–84.

16. Qutub AA, Ryan DT, Long B, Zaunbrecher R, Hu CW, Slater JH, et al., inventorsAutomated method for measuring, classifying, and matching the dynamics and information passing of single objects within one or more images2013.

17. Rekhi R, Ryan DT, Zaunbrecher R, Hu CW, Qutub AA. Computational Cell Phenotyping in the Lab, Plant, and Clinic. In: Zhang G, editor. Computational Bioengineering: CRC Press; 2015. p. 254–82.

18. Blankenship AG, Feller MB. Mechanisms underlying spontaneous patterned activity in developing neural circuits. Nature reviews Neuroscience. 2010;11(1):18–29.

19. Malmersjö S, Rebellato P, Smedler E, Planert H, Kanatani S, Liste I, et al. Neural progenitors organize in small-world networks to promote cell proliferation. Proceedings of the National Academy of Sciences of the United States of America. 2013;110:E1524–32.

20. Shimojo H, Ohtsuka T, Kageyama R. Oscillations in Notch Signaling Regulate Maintenance of Neural Progenitors. Neuron. 2008;58:52–64.

21. Otani T, Marchetto MC, Gage FH, Simons BD, Livesey FJ. 2D and 3D Stem Cell Models of Primate Cortical Development Identify Species-Specific Differences in Progenitor Behavior Contributing to Brain Size. Cell Stem Cell. 2016;18(4):467–80.

22. Li C, Xu D, Ye Q, Hong S, Jiang Y, Liu X, et al. Zika Virus Disrupts Neural Progenitor Development and Leads to Microcephaly in Mice. 2016.

23. Cai L, Hayes NL, Nowakowski RS. Synchrony of clonal cell proliferation and contiguity of clonally related cells: production of mosaicism in the ventricular zone of developing mouse neocortex. The Journal of neuroscience : the official journal of the Society for Neuroscience. 1997;17(6):2088–100.

24. Reznikov K, van der Kooy D. Variability and partial synchrony of the cell cycle in the germinal zone of the early embryonic cerebral cortex. The Journal of comparative neurology. 1995;360(3):536–54.

25. Sakaue-Sawano A, Kurokawa H, Morimura T, Hanyu A, Hama H, Osawa H, et al. Visualizing Spatiotemporal Dynamics of Multicellular Cell-Cycle Progression. Cell. 2008;132(3):487–98.

26. Megat S, Ray PR, Tavares-Ferreira D, Moy JK, Sankaranarayanan I, Wanghzou A, et al. Differences between Dorsal Root and Trigeminal Ganglion Nociceptors in Mice Revealed by Translational Profiling. J Neurosci. 2019;39(35):6829–47.

27. Kim YS, Anderson M, Park K, Zheng Q, Agarwal A, Gong C, et al. Coupled Activation of Primary Sensory Neurons Contributes to Chronic Pain. Neuron. 2016;91(5):1085–96.

28. Kim YS, Chu Y, Han L, Li M, Li Z, LaVinka PC, et al. Central terminal sensitization of TRPV1 by descending serotonergic facilitation modulates chronic pain. Neuron. 2014;81(4):873–87.

29. López-Cancio E, Ricciardi AC, Sobrino T, Cortés J, de la Ossa NP, Millán M, et al. Reported Prestroke Physical Activity Is Associated with Vascular Endothelial Growth Factor Expression and Good Outcomes after Stroke. Journal of Stroke and Cerebrovascular Diseases. 2017;26(2):425–30.

30. Wei ZZ, Zhang JY, Taylor TM, Gu X, Zhao Y, Wei L. Neuroprotective and regenerative roles of intranasal Wnt-3a Administration after focal ischemic stroke in mice. Journal of Cerebral Blood Flow & Metabolism. 2017:0271678X1770266–0271678X1770266.

31. Goodwin AM. In vitro assays of angiogenesis for assessment of angiogenic and anti-angiogenic agents. Microvascular Research. 2007;74(2):172–83.

32. Costa G, Harrington KI, Lovegrove HE, Page DJ, Chakravartula S, Bentley K, et al. Asymmetric division coordinates collective cell migration in angiogenesis. Nature Cell Biology. 2016;18(12):1292–301.

33. Slater JH, Culver JC, Long BL, Hu CW, Hu J, Birk TF, et al. Recapitulation and Modulation of the Cellular Architecture of a User-Chosen Cell of Interest Using Cell-Derived, Biomimetic Patterning. ACS Nano. 2015;9(6):6128–38.

34. Snijder B, Sacher R, Ramo P, Damm E-M, Liberali P, Pelkmans L. Population context determines cell-to-cell variability in endocytosis and virus infection. Nature. 2009;461(7263):520–3.

35. Gut G, Tadmor MD, Pe’er D, Pelkmans L, Liberali P. Trajectories of cell-cycle progression from fixed cell populations. Nature methods. 2015;12(10):951–4.

36. Vaicik MK, Thyboll Kortesmaa J, Moverare-Skrtic S, Kortesmaa J, Soininen R, Bergstrom G, et al. Laminin alpha4 deficient mice exhibit decreased capacity for adipose tissue expansion and weight gain. PLoS One. 2014;9(10):e109854.

37. Vaicik MK, Blagajcevic A, Ye H, Morse MC, Yang F, Goddi A, et al. The Absence of Laminin α4 in Male Mice Results in Enhanced Energy Expenditure and Increased Beige Subcutaneous Adipose Tissue. Endocrinology. 2018;159(1):356–67.

38. Vaicik MK, Morse M, Blagajcevic A, Rios J, Larson J, Yang F, et al. Hydrogel-Based Engineering of Beige Adipose Tissue. J Mater Chem B. 2015;3(40):7903–11.

39. Yang F, Cohen RN, Brey EM. Optimization of Co-Culture Conditions for a Human Vascularized Adipose Tissue Model. Bioengineering (Basel). 2020;7(3).

40. Berg S, Kutra D, Kroeger T, Straehle CN, Kausler BX, Haubold C, et al. ilastik: interactive machine learning for (bio)image analysis. Nature Methods. 2019;16(12):1226–32.

41. Hollandi R, Szkalisity A, Toth T, Tasnadi E, Molnar C, Mathe B, et al. nucleAIzer: A Parameter-free Deep Learning Framework for Nucleus Segmentation Using Image Style Transfer. Cell Systems. 2020;10(5):453–8.e6.

42. Moen E, Borba E, Miller G, Schwartz M, Bannon D, Koe N, et al. Accurate cell tracking and lineage construction in live-cell imaging experiments with deep learning. bioRxiv. 2019:803205–.

43. Carpenter AE, Jones TR, Lamprecht MR, Clarke C, Kang IH, Friman O, et al. CellProfiler: image analysis software for identifying and quantifying cell phenotypes. Genome Biology. 2006;7(10):R100–R.

44. Smedler E, Malmersjo S, Uhlen P. Network analysis of time-lapse microscopy recordings. Front Neural Circuits. 2014;8:111.

45. Bounova G, De Weck O. Overview of metrics and their correlation patterns for multiple-metric topology analysis on heterogeneous graph ensembles. Physical Review E - Statistical, Nonlinear, and Soft Matter Physics. 2012;85.

46. Lacoste A, Berenshteyn F, Brivanlou AH. An Efficient and Reversible Transposable System for Gene Delivery and Lineage-Specific Differentiation in Human Embryonic Stem Cells. Cell Stem Cell. 2009;5(3):332–42.

47. Ryan DT, Hu J, Long BL, Qutub AA. Predicting endothelial cell phenotypes in angiogenesis2013.

48. Hu CW, Li H, Qutub AA. Shrinkage Clustering: a fast and size-constrained clustering algorithm for biomedical applications. BMC Bioinformatics. 2018;19(1):19.

49. Snijder B, Sacher R, Rämö P, Liberali P, Mench K, Wolfrum N, et al. Single-cell analysis of population context advances RNAi screening at multiple levels. Molecular Systems Biology. 2012;8.

50. Schapiro D, Jackson HW, Raghuraman S, Fischer JR, Zanotelli VRT, Schulz D, et al. histoCAT: analysis of cell phenotypes and interactions in multiplex image cytometry data. Nature Methods. 2017.

51. Popovic D, Koch B, Kueblbeck M, Ellenberg J, Pelkmans L. Multivariate Control of Transcript to Protein Variability in Single Mammalian Cells. Cell Systems. 2018;7(4):398–411.e6.

52. Irmisch A, Bonilla X, Chevrier S, Lehmann KV, Singer F, Toussaint NC, et al. The Tumor Profiler Study: integrated, multi-omic, functional tumor profiling for clinical decision support. Cancer Cell. 2021.

53. Rose F, Rappez L, Triana SH, Alexandrov T, Genovesio A. PySpacell: A Python Package for Spatial Analysis of Cell Images. Cytometry Part A. 2019.

54. Svensson V, Teichmann SA, Stegle O. SpatialDE: Identification of spatially variable genes. Nature Methods. 2018;15(5):343–6.

55. Edsgärd D, Johnsson P, Sandberg R. Identification of spatial expression trends in single-cell gene expression data. Nature Methods. 2018;15(5):339–42.

56. Stoltzfus CR, Filipek J, Gern BH, Olin BE, Leal JM, Wu Y, et al. CytoMAP: A Spatial Analysis Toolbox Reveals Features of Myeloid Cell Organization in Lymphoid Tissues. Cell Reports. 2020;31(3).

57. Qin Y, Winsnes CF, Huttlin EL, Zheng F, Ouyang W, Park J, et al. Mapping cell structure across scales by fusing protein images and interactions. bioRxiv. 2020:2020.06.21.163709–2020.06.21.

58. Romano SA, Pérez-Schuster V, Jouary A, Boulanger-Weill J, Candeo A, Pietri T, et al. An integrated calcium imaging processing toolbox for the analysis of neuronal population dynamics. PLOS Computational Biology. 2017;13(6):e1005526–e.

59. Giovannucci A, Friedrich J, Gunn P, Kalfon J, Brown BL, Koay SA, et al. CaImAn an open source tool for scalable calcium imaging data analysis. eLife. 2019;8.

60. Cantu DA, Wang B, Gongwer MW, He CX, Goel A, Suresh A, et al. EZcalcium: Open Source Toolbox for Analysis of Calcium Imaging Data. bioRxiv. 2020:2020.01.02.893198–2020.01.02.

61. Prada J, Sasi M, Martin C, Jablonka S, Dandekar T, Blum R. An open source tool for automatic spatiotemporal assessment of calcium transients and local ‘signal-close-to-noise’ activity in calcium imaging data. PLOS Computational Biology. 2018;14(3):e1006054–e.

62. Tegtmeier J, Brosch M, Janitzky K, Heinze H-J, Ohl FW, Lippert MT. CAVE: An Open-Source Tool for Combined Analysis of Head-Mounted Calcium Imaging and Behavior in MATLAB. Frontiers in Neuroscience. 2018;12:958–.

63. Moein M, Grzyb K, Gonçalves Martins T, Komoto S, Peri F, Crawford AD, et al. CaSiAn: a Calcium Signaling Analyzer tool. Bioinformatics. 2018;34(17):3052–4.

64. Kaifosh P, Zaremba JD, Danielson NB, Losonczy A. SIMA: Python software for analysis of dynamic fluorescence imaging data. Frontiers in Neuroinformatics. 2014;8.

65. Pachitariu M, Stringer C, Schröder S, Dipoppa M, Rossi LF, Carandini M, et al. Suite2p: beyond 10,000 neurons with standard two-photon microscopy. bioRxiv. 2016:061507–.

66. Zhou P, Resendez SL, Rodriguez-Romaguera J, Jimenez JC, Neufeld SQ, Giovannucci A, et al. Efficient and accurate extraction of in vivo calcium signals from microendoscopic video data. eLife. 2018;7.

67. Reynolds S, Abrahamsson T, Schuck R, Sjöström PJ, Schultz SR, Dragotti PL. ABLE: an Activity-Based Level Set Segmentation Algorithm for Two-Photon Calcium Imaging Data. bioRxiv. 2017:190348–.

68. Petersen A, Simon N, Witten D. Scalpel: Extracting neurons from calcium imaging data. Annals of Applied Statistics. 2018;12(4):2430–56.

69. Lu J, Li C, Singh-Alvarado J, Zhou ZC, Fröhlich F, Mooney R, et al. MIN1PIPE: A Miniscope 1-Photon-Based Calcium Imaging Signal Extraction Pipeline. Cell Reports. 2018;23(12):3673–84.

70. Rueckl M, Lenzi SC, Moreno-Velasquez L, Parthier D, Schmitz D, Ruediger S, et al. SamuROI, a Python-Based Software Tool for Visualization and Analysis of Dynamic Time Series Imaging at Multiple Spatial Scales. Frontiers in Neuroinformatics. 2017;11.

71. Colizza V, Flammini A, Serrano MA, Vespignani A. Detecting rich-club ordering in complex networks. 2006;2(February):110–5.

72. Newman M. Assortative Mixing in Networks. Physical Review Letters. 2002;89(20):208701–.

