## Supplementary Information for "cytoNet: Spatiotemporal Network Analysis of Cell Communities"

| | |
|---|---|
| **Supplementary Figure 1** | Image segmentation of HUVEC immunofluorescence images |
| **Supplementary Figure 2** | Image segmentation of FUCCI-ReN nucleus images |
| **Supplementary Figure 3** | Correlation heatmap of local network metrics and morphology metrics for immunofluorescence HUVEC images |
| **Supplementary Table 1** | Metrics used to define endothelial cell morphology |
| **Supplementary Table 2** | Cohen's d effect size for treatment conditions on morphology metrics shown in Figure 3 (f-h) in the main text |
| **Supplementary Video 1**<br>Video 1 link | Time-lapse movie of sparse culture of FUCCI-ReN cells |
| **Supplementary Video 2**<br>Video 2 link | Time-lapse movie of sparse culture of FUCCI-ReN cells, with cell boundaries and graph representation overlaid |
| **Supplementary Video 3**<br>Video 3 link | Time-lapse movie of dense culture of FUCCI-ReN cells |
| **Supplementary Video 4**<br>Video 4 link | Time-lapse movie of dense culture of FUCCI-ReN cells, with cell boundaries and graph representation overlaid |
| **Supplementary Methods 1** | Instructions for using the cytoNet web-based user interface |

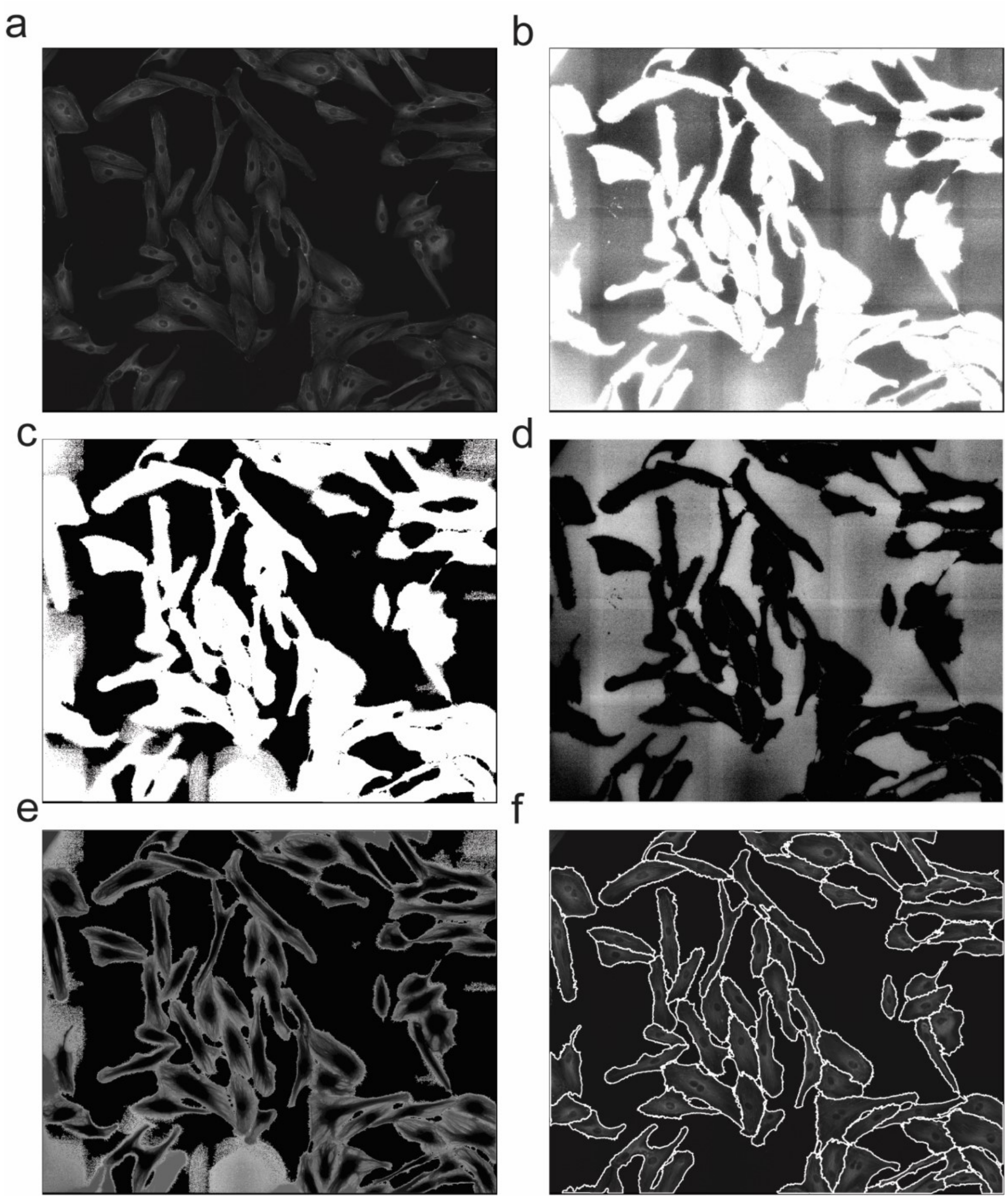


**Supplementary Figure 1. Image segmentation of HUVEC immunofluorescence images. (a)** Original grayscale image. **(b)** Image after adaptive histogram equalization and Gaussian filtering. **(c)** Binary image obtained using Otsu's threshold, with small objects removed. **(d)** Complement of filtered image in **(b)**. **(e)** Watershed basins obtained through imposing minimum of images in **(d)** and the marker image (obtained by combining the binary image in **(c)** and the image obtained through binarization of microtubules and nuclei). **(f)** Final cell borders.

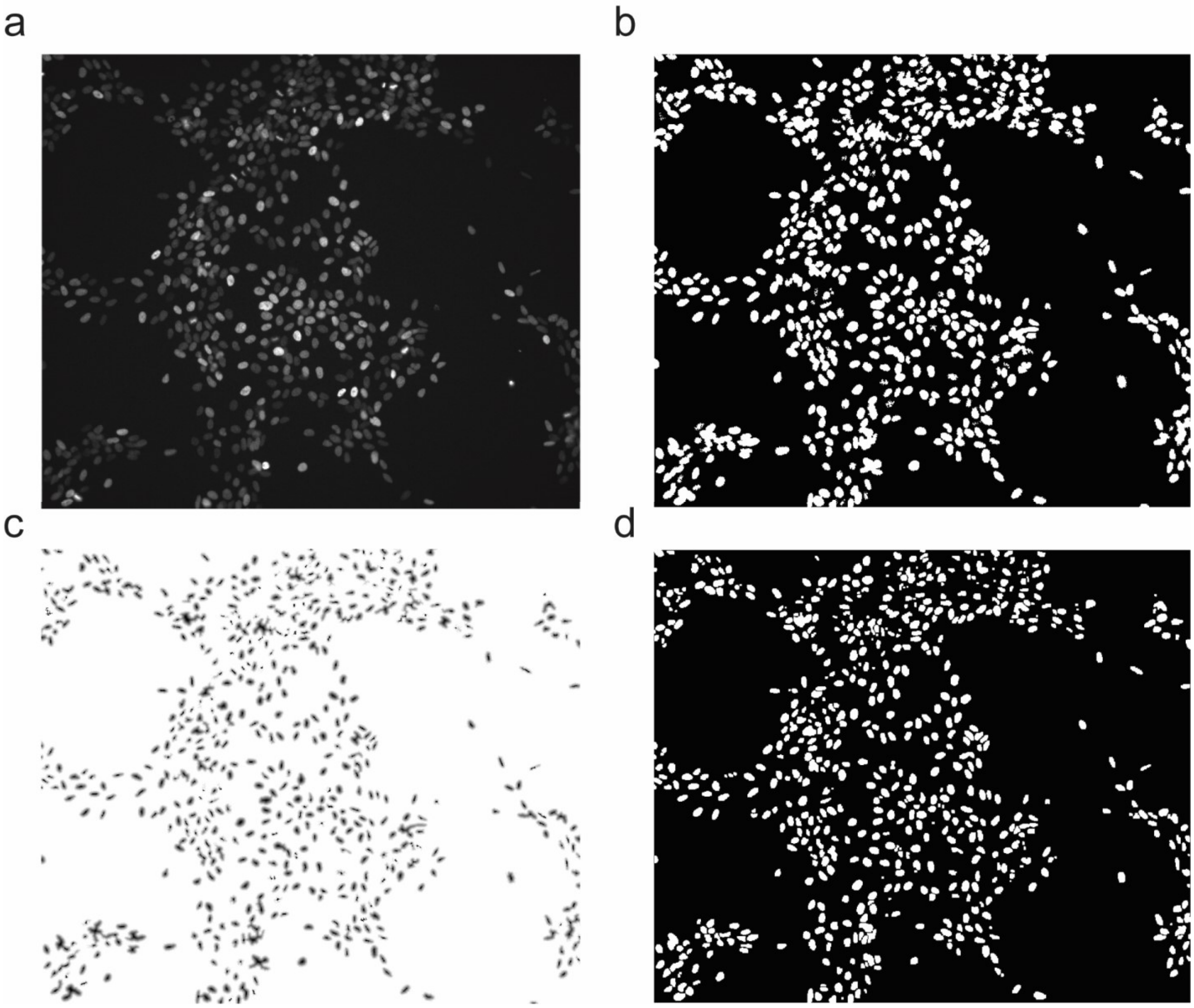


**Supplementary Figure 2. Image processing steps for FUCCI-ReN nucleus images. (a)** Fluorescence image from H2B-Cerulean channel marking all nuclei. **(b)** Binary mask obtained through adaptive thresholding. **(c)** Image obtained through imposing minimum of distance transform of binary image in **(b)** and local minima. This image serves as seeds for the watershed algorithm. **(d)** Final mask obtained after watershed transform.

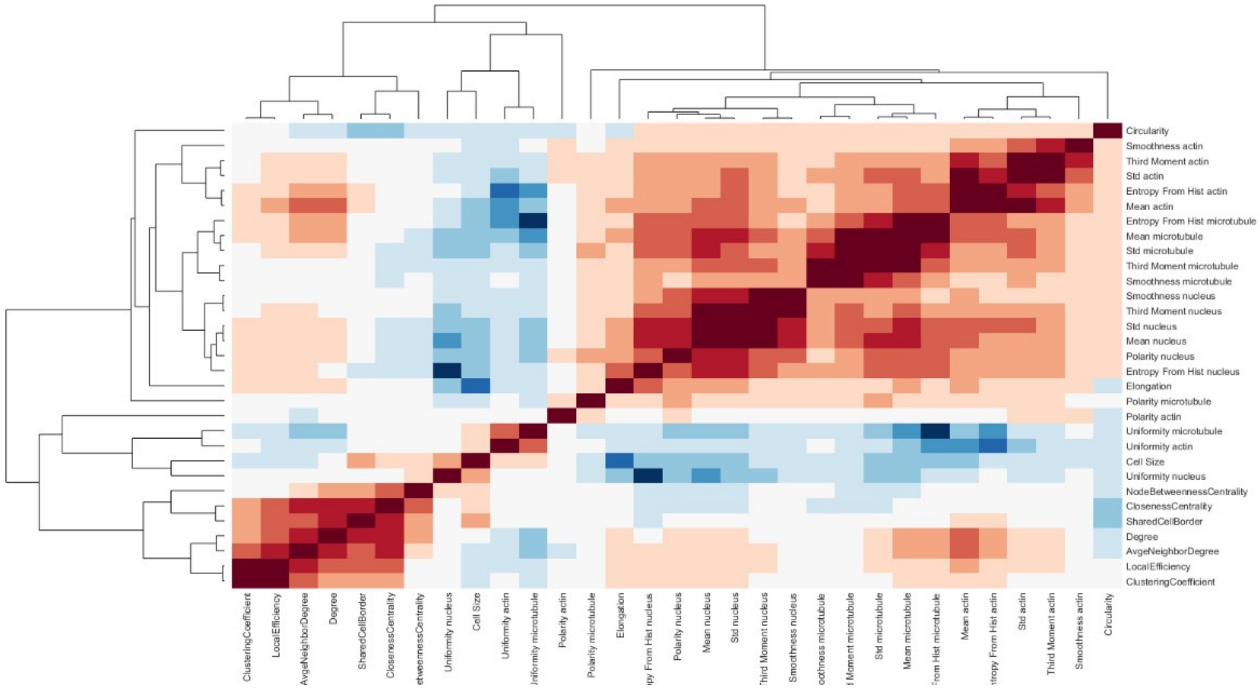


**Supplementary Figure 3. Correlation heatmap of local network metrics and morphology metrics for immunofluorescence HUVEC images**. All morphology and local network metrics (**Supplementary Table 1, Supplementary Table 2**) were combined into a single matrix. The cluster dendrogram was obtained through hierarchical clustering of the covariance matrix using Pearson's correlation as the similarity metric.

**Supplementary Table 1. Metrics used to define endothelial cell morphology.**

| Metrics | Definition | Mathematical Representation |
|---|---|---|
| Cell Size | Cell Spread Area | $A_c$ |
| Circularity | Shape factor | $\frac{4\pi A_c}{P_c}$ Where $P_c$ is perimeter of cell |
| Elongation | Shape factor | $\frac{P_c}{A_c}$ |
| Polarity* | Distance between center of mass of stain and the centroid of the cell | $\sqrt{(\chi_{C,x} - \Omega_{S,x})^2 + (\chi_{C,x} + \Omega_{S,x})^2}$ Where $\chi_c$ is centroid of cell and $\Omega_S$ is center of mass of stain |
| Mean* | First moment of grayscale stain intensity distribution | $\sum_{i=0}^{255} \frac{i}{255} \cdot p$ Where $p$ is the histogram counts of the image for pixel intensities, with 256 possible bins for a grayscale image |
| Standard Deviation* | Second moment of grayscale stain intensity distribution | $\sqrt{\sum_{i=0}^{255} \left(\frac{i}{255}\right)^2 \cdot p}$ |
| Third Moment* | Third moment of grayscale stain intensity distribution | $\frac{1}{255^2} \sqrt{\sum_{i=0}^{255} \left(\frac{i}{255}\right)^3 \cdot p}$ |
| Smoothness* | Measure of smoothness of stain | $1 - \frac{1}{1 + \left(\frac{1}{255^2} \sqrt{\sum_{i=0}^{255} \left(\frac{i}{255}\right)^2 \cdot p}\right)}$ |
| Entropy from Histogram* | Measure of randomness of the stain intensity | $-\sum p \cdot log_2(p)$ |
| Uniformity* | Sum of squared elements in the histogram counts of the image for pixel intensities | $\sum p^2$ |

*Computed for all 3 stain

**Supplementary Table 2. Cohen's d effect size for treatment conditions on morphology metrics shown in Figure 4(g-h) in the main text.**

| Morphology Metric | Treatment Condition | Cohen's d Effect Size | | | |
|---|---|---|---|---|---|
| | | No correction for network metrics | | Correction applied for network metrics | |
| Cell.Size | | 6hr (uncorrected*) | 12hr (uncorrected) | 6hr (corrected**) | 12hr (corrected) |
| | BDNF50 | 0.256 | 0.217 | 0.148 | 0.170 |
| | VEGF50 | 0.151 | 0.023 | 0.093 | 0.068 |
| Mean.Actin | | 6hr (uncorrected) | 12hr (uncorrected) | 6hr (corrected) | 12hr (corrected) |
| | BDNF50 | 0.381 | 1.020 | 0.091 | 0.873 |
| | BDNF100 | 0.517 | 2.522 | 0.260 | 1.959 |
| | VEGF50 | 1.121 | 1.018 | 0.348 | 0.740 |
| | VEGF100 | 1.267 | 2.269 | 0.284 | 1.808 |

**Supplementary Video 1**

Time-lapse movie of sparse culture of FUCCI-ReN cells. Magenta: Cdt1-mCherry, Green: Geminin-Venus. Time stamp is shown on top right corner.

**Supplementary Video 2**

Time-lapse movie of sparse culture of FUCCI-ReN cells with graph overlay. Movie displays phase contrast frames from movie in Supplementary Video 1, with Cdt1(-)/mCherry(+) nuclei circled in magenta, Geminin(-)/Venus(+) nuclei circled in green and mCherry(-)/Venus(-) nuclei circled in blue. Yellow lines represents proximity edges.

**Supplementary Video 3**

Time-lapse movie of dense culture of FUCCI-ReN cells. Magenta: Cdt1-mCherry, Green: Geminin-Venus. Time stamp is shown on top right corner.

**Supplementary Video 4**

Time-lapse movie of dense culture of FUCCI-ReN cells with graph overlay. Movie displays phase contrast frames from movie in Supplementary Video 3, with Cdt1(-)/mCherry(+) nuclei circled in magenta, Geminin(-)/Venus(+) nuclei circled in green and mCherry(-)/Venus(-) nuclei circled in blue. Yellow lines represents proximity edges.

**Supplementary Methods 1**

**Instructions for using the web-based user interface**

Go to https://www.qutublab.org/how and follow the "cytoNet site" link. An explanation of parameters and input format can also be downloaded there.

1. Select mask files

a) Select binary mask files by clicking on the 'Browse' button to start a file selection dialog box. Mask file names must start with the prefix "MASK_". Multiple files can be selected by: i) clicking on a file while holding down the control key (command key in MacOS; ii) clicking and dragging; or iii) entering control-a (command-a in MacOS) to select all files in a directory or folder.

b) For demonstration purposes, cytoNet can provide a mask image if you do not have your own. Check the box next to the sample mask to include it as input.

2. Select edge determination method for spatial graphs

a) Cell Centroid Distance. Edges between nearby objects are determined by the distance between their centroids. If this method is selected, use the slider bar to specify an adjacency threshold. The adjacency threshold determines the maximum distance between two centroids at which an edge is created in the following way. Let $a_1$ and $a_2$ be the area of two objects with centroids $c_1$ and $c_2$ respectively. For each object, compute its effective radius: $r_i = \sqrt{a_i/\pi}$. A graph edge is placed between two objects (vertices) whenever the distance between their centroids is within the adjusted sum of their effective radii:

$distance(c_1, c_2) \leq S \cdot (r_1 + r_2)$ where $S$ is the user defined adjacency threshold parameter.

b) Border Overlap. Edges between objects are determined by the sharing of border pixels.

c) Cell Perimeter Distance. Edges between nearby objects are determined by the shortest distance between pixels of the two objects. If this method is selected, specify the maximum distance in pixels between objects that can be connected by an edge.

3. Select image files (optional)

a) Select image files by clicking on the ‘Browse’ button to start a file selection dialog box. Image file names must match mask file names after the "MASK_" prefix is removed. Multiple files can be selected in the same manner as for mask files. Color input images are first converted to grayscale images by cytoNet before being processed as previously described.

b) If the image files contain sequences of calcium signal images, check the box indicating calcium signaling.

c) For demonstration purposes, cytoNet can provide a mask image if you do not have your own. Check the box next to the sample image to include it as input.

4. Enter an email address. cytoNet will use this email address to inform you that processing is complete.

5. Click the Submit button.

6. cytoNet will send you an email message indicating that your request has been accepted. This message includes a Request ID that you can use to check on the progress of your request. cytoNet will also send you an email message informing you that processing has ended for your request.

7. When your request has been successfully processed, you may download your results. Note that your results will be available for only a limited amount of time.

Results are formatted as follows. Global metrics are tabulated in a file called ‘GlobalMetrics.csv’ for all images in the input folder. Local metrics, computed on a per-cell basis are tabulated in a separate file for each image called ‘LocalMetrics_filename.csv’, where filename is the original file name. Also, basic morphology metrics (size, elongation, circularity and stain intensity) are tabulated in a separate file for each image called SingleCellMetrics_filename.csv, where filename is the original file name. Processed images are also created for each image in the input folder, called ‘filename_processed.tif’ where the original image is overlaid with cell indices, object outlines (red) and spatial proximity edges (yellow). Cell indices displayed in the processed images are used in the first column of local metrics and single cell metric files.
